# Pulmonary Hypertension Promotes Neuroinflammation and Neurodegeneration

**DOI:** 10.1101/2025.09.02.673803

**Authors:** Priscilla M. De La Cruz, Angelia Lockett, Marta T. Gomes, Somanshu Banerjee, Asif Razee, Amanda Fisher, Todd Cook, Christopher D. Lloyd, Shino Magaki, Soban Umar, Adrian L. Oblak, Roberto F. Machado

## Abstract

**Introduction:** Pulmonary arterial hypertension (PAH) is associated with neurocognitive deficits and abnormal brain MRI. Little is known about the mechanisms underlying these clinical observations. TDP-43 is a proteinopathy associated with frontotemporal lobar degeneration (FTLD), Alzheimer’s Disease, and Amyotrophic Lateral Sclerosis (ALS). In this study, we hypothesize PAH will result in gliosis, reduced neuronal density, and increased TDP-43 mislocalization.

**Methods:** Sprague Dawley rats were randomly assigned to receive Vehicle (DMSO) Monocrotaline, or Sugen/Hypoxia to induce PH. Right heart catheterization was used to confirm PAH. Brain tissue was fixed and probed for microglia (Iba1), astrocytes (GFAP), neurons (NeuN), and TDP-43. Human PH vs control brain tissue was also probed for NeuN and TDP-43.

**Results and Conclusions:** We identify an increase in microglia and astrocyte density in the frontal cortex along with reduced neuronal density and neuronal TDP-43 mislocalization in rat models of PH. In addition, human PH frontal cortex demonstrated neuronal TDP-43 mislocalization. This is the first evidence of TDP-43 proteinopathy in PH.

## Introduction

Pulmonary hypertension (PH) is a severe condition, defined by an elevated mean pulmonary arterial pressure exceeding 20 mmHg, and affects approximately 1% of the global population^1^. Clinical studies indicate that patients with PH exhibit cognitive impairments, including challenges in executive function, memory, calculation, and visuospatial abilities^2–4^. A small imaging study reveals a reduction in grey matter volume on brain magnetic resonance imaging when compared to healthy age-matched control subjects, indicating potential neuronal loss ^5^. Research on the pathophysiological effects of PH on the central nervous system remains limited. Animal studies have indicated increased concentrations of microglia and astrocytes in the hypothalamus and thoracic spinal cord in subjects with PH compared to control groups, which may correlate with the severity of PH^6,7^. The integration of clinical and animal data suggests the presence of neuroinflammatory and neurodegenerative processes.

TAR DNA-binding protein 43 (TDP-43), a critical regulator of RNA metabolism, has been extensively implicated in neurodegenerative disorders such as amyotrophic lateral sclerosis (ALS) and frontotemporal lobar degeneration (FTLD)^8–10^. Pathological mislocalization and aggregation of TDP-43 disrupt RNA splicing, translation, and intracellular transport—processes essential to both neuronal and endothelial cell function^11,12^. Recent data suggest a potential link between TDP-43 and inflammation, implicating it as a convergent node in pulmonary hypertension (PH)-associated multi-organ dysfunction. Notably, pathologic extranuclear TDP-43 aggregation occurs under pro-inflammatory conditions such as stroke and traumatic brain injury^11,12^. The role of TDP-43 in PH-associated neurodegeneration has not yet been explored.

In this study, we investigated the role of pulmonary hypertension (PH) in cortical neuroinflammation and neurodegeneration through complementary methodologies. We characterized brain tissue from animal models of PH to evaluate astrocyte, microglia, and neuronal densities, as well as TDP-43 proteinopathy. Additionally, we examined neuronal density and TDP-43 proteinopathy in brain tissue samples from PH patients. Given the emerging evidence linking TDP-43 to endothelial dysfunction, we conducted a secondary analysis of pulmonary artery endothelial cell (PAEC) transcriptomic data to identify potential TDP-43-regulated genes. While our studies on animal and human tissues concentrated on central nervous system (CNS) pathology, these transcriptomic insights suggest a vascular component to the neuroinflammatory processes observed in PH. We hypothesize that PH induces cortical neuroinflammation, neurodegeneration, and TDP-43 proteinopathy.

## Methods

### Animal Models of Pulmonary Hypertension and Hemodynamic Measurements

All experimental procedures were sanctioned by the Institutional Animal Care and Use Committee at Indiana University. Following standard protocol as previously described^13^, male and female Sprague Dawley rats (190-200 g) were procured from Charles River and randomly allocated to receive either Vehicle (Dimethyl sulfoxide (DMSO), intraperitoneal (ip) vehicle injection as a control), Monocrotaline (MCT; ip, 60 mg/kg of body weight), or Sugen (subcutaneously, 20 mg/kg body weight). Rats administered Sugen were housed in a 10% oxygen chamber for three weeks, followed by two weeks under normoxia conditions. At five weeks post-injection, the rats underwent open field testing, as previously described^14^. Briefly, the rats were placed in an open field maze, and measurements of distance traveled, time spent in the center, time at rest, and average ambulatory velocity were recorded using Any-maze software. Right Ventricular Systolic Pressure (RVP) was assessed via right heart catheterization employing a Millar pressure transducer catheter at the time of euthanasia. The Fulton index was calculated by measuring the ratio of the right ventricle weight to the sum of the weights of the left ventricle and septum (RV/(LV+S)). Brain tissue and plasma were collected at the time of euthanasia.

### Rat Brain Tissue Immunofluorescence Staining

The brains were meticulously dissected along the intracerebral fissure, with one half snap-frozen in liquid nitrogen and stored at -80°C for subsequent RNA analyses. The remaining half was allocated for immunofluorescence studies and was fixed with 4% paraformaldehyde overnight, followed by a 24-hour treatment with 30% sucrose. The brains were then stored in optimal tissue cutting medium at -80°C until sectioning. Coronal sections were prepared at a thickness of 30 micrometers and preserved in cryoprotectant solution (FD NeuroTechnologies, Inch, Cat#PC102). The sections underwent heat-mediated antigen retrieval for 30 minutes at 80°C (Universal HEIR antigen retrieval reagent, Abcam#ab208572). Subsequently, the sections were washed with 0.3% Phosphate Buffered Saline-Triton X-100 (PBST) and subjected to a 10-minute treatment with 0.3% Hydrogen Peroxide block. Following additional washing, the sections were incubated with 10% Donkey Serum (Abcam, #NC1697010) for 1 hour at room temperature. The sections were then incubated overnight with primary antibodies (Supplemental Table 1). After the overnight incubation, the sections were washed with PBST, blocked with 10% Donkey Serum, and incubated with secondary antibodies (1:1000) for 2 hours. The sections were then washed with PBST and mounted on slides using ProLong Gold Antifade Mountant with DNA Stain DAPI (Invitrogen, #P36931). Imaging was conducted using the Leica Thunder System (20x magnification unless specified), and the images were analyzed using Imaris (Oxford Instruments) to quantify cells positively expressing both DAPI and cell-specific marker.

### Rat Brain Quantitative Polymerase Chain Reaction (PCR)

RNA was extracted from hemibrains utilizing the RNeasy Plus Mini Kit (Qiagen, # 74134) in accordance with the manufacturer’s protocol. The quality of the RNA was evaluated using the NanoDrop One. Subsequently, RNA was transcribed into cDNA employing the High-Capacity cDNA Reverse Transcription Kit (Applied Biosystems, # 4368814) with the Bio-rad T100 Thermal Cycler. Relative quantitative RT-PCR was conducted using the Roche Lightcycler 480 and the PowerUP SYBR Green Master Mix chemistry (Applied Biosystems, # A25742) to quantify cytokine gene expression. The primers used were as follows: *IL-6* (Forward ‘TCCTACCCCAACTTCCAATGCTC’, Reverse ‘TTGGATGGTCTTGGTCCTTAGCC’), *IL-1β* (Forward ‘CACCTCTCAAGCAGAGCACAG’, Reverse ‘GGGTTCCATGGTGAAGTCAAC’), *TNF-α* (Forward ‘AAATGGGCTCCCTCTCATCAGTTC’, Reverse ‘TCTGCTTGGTGGTTTGCTACGAC’), and *Gapdh* (Forward ‘CATCACTGCCACCCAGAAGACTG’, Reverse ‘ATGCCAGTGAGCTTCCCGTTCAG’). The fold change was calculated using the 2^-ΔΔCT^ method, with results normalized to *Gapdh*.

### Human Cortical Tissue Immunohistochemistry

In accordance with 45 CFR 46, our study does not involve human subjects but used deidentified human tissues provided by UCLA Pathology departmental honest broker, the UCLA Translational Pathology Core Lab (IRB# 11-002504). Paraffin-embedded 5 micrometer thick human brain cortex sections (Control n=3, PH n=4) were rehydrated, and antigen retrieval was performed by heat-induced epitope retrieval at pH 6 (C9999, Sigma). Tissue sections were blocked for an hour at room temperature in PBS with 5% goat serum, 1% BSA, 0.1% Triton, 0.05% Tween, and 0.3 M glycine. Tissue sections were incubated overnight with primary antibodies diluted 1:200 in PBS with 1% BSA and 0.5% Triton at 4°C. Secondary antibodies were incubated for an hour at room temperature and diluted 1:1000 in PBS with 0.05% Tween. Sections were incubated with ProLong™ Gold antifade mountant with DNA Stain DAPI and cover slips were applied. All images were acquired using a confocal microscope (40x magnification) (Nikon). A total of 4-5 images were acquired from each slide. Images were analyzed using Imaris (Oxford Instruments).

### Secondary Analysis of TDP-43-Regulated Genes in Pulmonary Artery Endothelial Cells

Raw gene expression count data were procured from the NCBI Gene Expression Omnibus (GEO) under accession number GSE243193, encompassing RNA-sequencing data from human pulmonary artery endothelial cells (PAECs) isolated from patients with group 1 pulmonary hypertension (PH) (n=20) and controls (n=3) ^15^. Group 1 PH patients were selected as they more closely align with the animal models represented in this study. The dataset was imported into R (version 4.2.0) and underwent preprocessing to eliminate genes with missing or NA values across samples^16^. Following quality control, expression counts were converted into a matrix. Samples were categorized as “PH” or “Control” based on their column names, and group labels were encoded as a factor variable for subsequent differential expression analysis. Gene expression normalization and differential analysis were conducted using the edgeR and limma packages^17,18^. Expression counts were filtered using filterByExpr() to retain genes with adequate counts across samples. Library sizes were normalized using the trimmed mean of M-values (TMM), and log2-transformed counts per million (CPM) were calculated via voom to model the mean-variance relationship^19^. Linear modeling was executed using lmFit(), followed by empirical Bayes moderation using eBayes() to enhance variance estimation. Differentially expressed genes (DEGs) were identified based on false discovery rate (FDR)-adjusted p-values < 0.05. Gene annotation was performed using the org.Hs.eg.db Bioconductor package to map ENTREZ identifiers to gene symbols and gene names^20^. To evaluate the enrichment of TDP-43 regulatory activity in PH-associated gene expression changes, a curated list of TDP-43 RNA targets was obtained from POSTAR3 CLIP-seq datasets^21^. POSTAR-3 utilizes data from the ENCODE project which identified TDP-43 targets under baseline conditions. Although these targets were identified in non-diseased immortalized cells, they provide a reference map of physiologic TDP-43 binding sites, which may be dysregulated or lost in disease-specific contexts such as nuclear exclusion^22^.The overlap between differentially expressed genes (FDR < 0.05) and TDP-43 targets was calculated. A Fisher’s exact test was conducted to assess whether TDP-43 targets were statistically overrepresented among DEGs compared to the background set of all expressed genes. A 2×2 contingency table was constructed with counts of genes categorized as TDP-43 target vs. non-target and DEG vs. non-DEG. The resulting odds ratio and p-value were utilized to determine enrichment significance. A volcano plot was generated using ggplot2, with upregulated and downregulated genes highlighted^23^. ChatGPT was used for code debugging.

### Statistical Analysis

Statistical analyses were performed utilizing GraphPad Prism (version 8.4.3; GraphPad Software) and R (version 4.2.0), as delineated in the preceding section. To compare experimental groups (e.g., SuHx vs. Vehicle and MCT vs. Vehicle), an unpaired two-tailed Student’s t-test was employed. Linear regression analysis was conducted to assess correlations between variables. Pearson’s correlation coefficient (r) and the coefficient of determination (R²) were calculated and reported. Data are presented as mean ± standard error of the mean (SEM) unless otherwise specified. A p-value < 0.05 was considered indicative of statistical significance.

## Results

### SuHx-Induced Pulmonary Hypertension is Associated with Elevated RVP and Hypertrophy

Sugen/hypoxia (SuHx) and monocrotaline (MCT) are established models for the study of pulmonary hypertension (PH) and were employed to induce PH in rats^24^. The presence of pulmonary hypertension was confirmed through measurements of right ventricular pressure (RVP) and right ventricular hypertrophy (Fulton Index). The SuHx model (n=6) elicited a pronounced PH phenotype, evidenced by a significant increase in RVP (mean difference of 52.33 ± 11.84 mmHg compared to vehicle controls, n=6; p=0.0013) and the Fulton index (mean difference of 0.2428 ± 0.03486, p<0.0001) (**Figure 1**). Conversely, the MCT model (n=6) demonstrated a trend towards increased RVP (mean difference of 17.33 ± 6.98 mmHg versus controls, p=0.0522) and a modest yet statistically significant increase in the Fulton Index (mean difference of 0.09815 ± 0.03066 versus controls, p=0.0198) (**Supplemental Figure 1**). Although this study was not designed or powered to assess sex differences (n=3 males and n=3 females per group), exploratory analysis indicated a sex-dependent effect in both models. In SuHx-treated rats, males exhibited higher RVP (mean: 102.67 mmHg) and Fulton Index (mean: 0.53) compared to females (mean RVP: 52 mmHg; mean Fulton index: 0.41). A similar pattern was observed in MCT-treated animals, with males showing higher RVP (mean: 56 mmHg) and Fulton Index (mean: 0.36) compared to females (mean RVP of 27 mmHg; mean Fulton index of 0.28). These findings align with existing literature on sex differences in experimental PH and highlight the necessity of considering sex as a biological variable in preclinical studies^25^. No statistically significant sex differences were observed in any of the subsequent measurements. This suggests a need for further exploration of the impact of sex as a biological variable in neuroinflammation in PH. Due to the relatively mild phenotype induced by MCT in our study (likely due to the modest effect on female mice), all MCT-associated findings, including cardiovascular and central nervous system assessments, are included in the supplemental materials.

**Figure 1.**
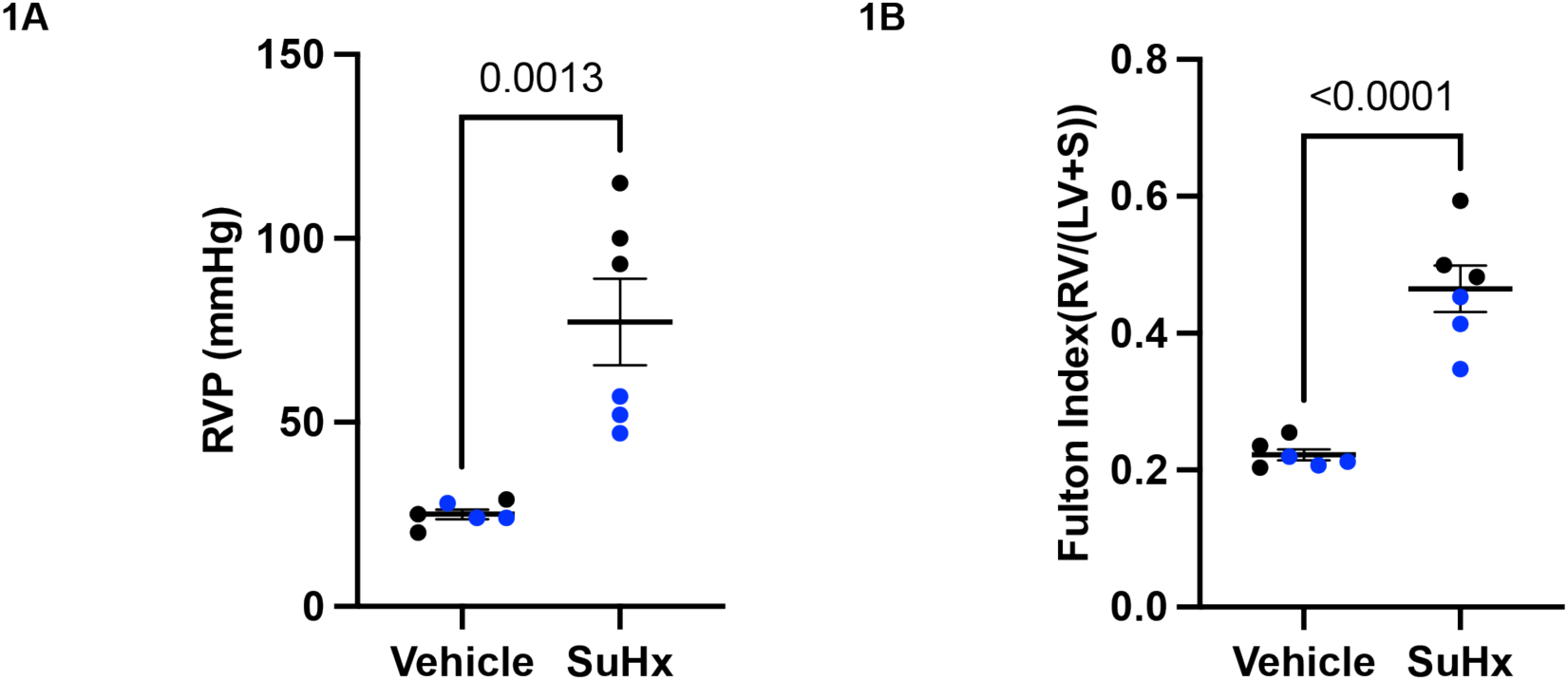
Sugen-hypoxia induces pulmonary hypertension and vascular remodeling. Rats treated with Sugen-hypoxia (SuHx; n = 6) exhibit increased right ventricular pressure (RVP), with a mean difference of 52.33 ± 11.84 mmHg compared to vehicle controls (n = 6; p = 0.0013) (**1A**). SuHx-treated rats also demonstrate right ventricular hypertrophy, with a mean Fulton Index difference of 0.2428 ± 0.03486 compared to controls (p < 0.0001) **(1B).** Data analyzed using a two-tailed unpaired t-test and are presented as mean ± SEM. Female animals are depicted in blue.

### No Behavioral Abnormalities are Observed in Open Field Testing

Open field testing (OFT) is widely used to assess sickness behavior, locomotor activity, and anxiety-like behavior in rodent models of systemic disease, including those of sepsis-associated encephalopathy, heart failure, and frontotemporal lobar degeneration^26–28^. In this study, OFT was employed to evaluate whether PH was associated with changes in exploratory behavior or activity levels. SuHx-treated rats showed no significant differences in total distance traveled, time spent in the center of the arena, time spent at rest, or average ambulatory velocity compared to vehicle-treated controls (**Figure 2**). Similarly, no statistically significant differences were observed between MCT-treated rats and controls (**Supplemental Figure 2**). OFT data did not correlate significantly with RVP or Fulton in either model (data not shown).

**Figure 2.**
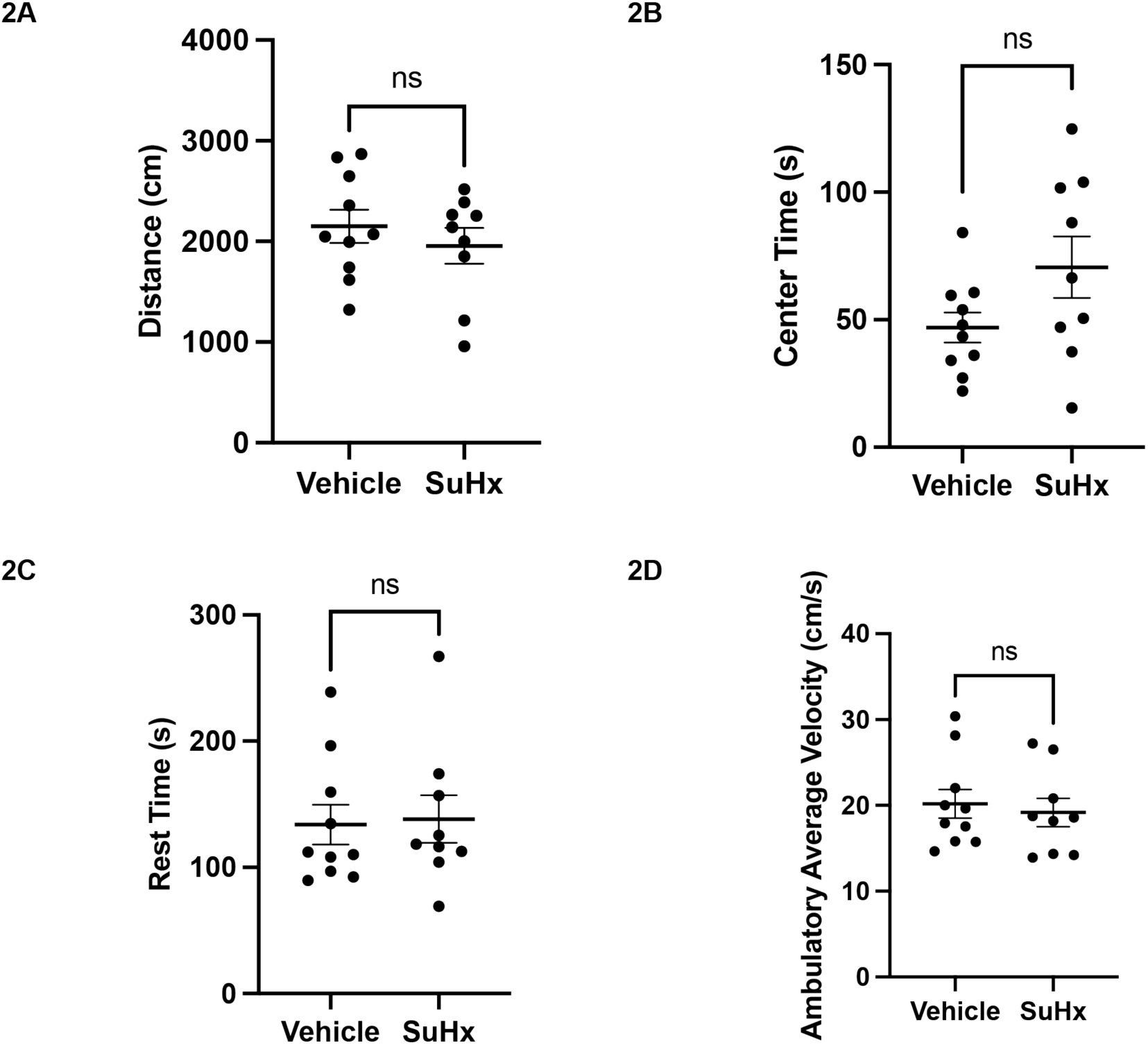
Sugen hypoxia does not induce sickness behavior in rats undergoing open field test. Rats exposed to Sugen hypoxia (SuHx, n = 9) did not exhibit significant differences in behavioral parameters compared to vehicle-treated controls (n = 10). Specifically, there were no significant changes in total distance traveled (**2A**), time spent in the center of the open field test (**2B**), time spent at rest (**2C**), or average ambulatory velocity (**2D**). Data were analyzed using an unpaired two-tailed t-test and are presented as mean ± SEM.

### Experimental PH Induces Neuroinflammation

Neuroinflammation is recognized as an early and critical risk factor for neurodegeneration^29,30^. In this study, we measured cortical density of microglia and astrocytes in two rodent models of PH. In SuHx-treated rats, microglial density was significantly increased (3.875 × 10^-5^ ± 1.504 × 10^-5^ cells/μm^2^ compared to vehicle-treated controls (n = 6; p = 0.0276). Astrocyte density was also elevated in SuHx animals (4.487 × 10^-5^± 1.806 × 10^-5^ cells/μm^2^, p = 0.0323) (**Figure 3**). Notably, neither microglia nor astrocytic density correlated with RVP or Fulton Index in the SuHx group (data not shown). In the MCT-treated rats, a significant increase in microglial density (0.2320 ± 0.009541 cells/μm^2^) compared to vehicle-treated controls (n = 6; p <0.0001) was observed (**Supplemental Figure 3**). Similarly, astrocyte density was elevated in MCT-treated animals relative to control (4.822 x 10^-5^ ± 1.309 x 10^-5^ cells/μm^2^, p= 0.0043) (**Supplemental Figure 3**). Unlike the SuHx-treated animals, cortical microglia density positively correlated with the RVP (r= 0.7670, 95% CI: 0.3449 to 0.9311, p=0.0036) and Fulton Index (r= 0.7990, 95% CI: 0.4156 to 0.9413, p=0.0018). Despite increased glial density, there was no increase in cortical expression of pro-inflammatory cytokines—including IL-1β, TNF-α, or IL-6 mRNA— between SuHx- or MCT-treated animals and their respective controls, with SuHx data presented in **Figure 4** and MCT data presented in **Supplemental Figure 4.**

**Figure 3.**
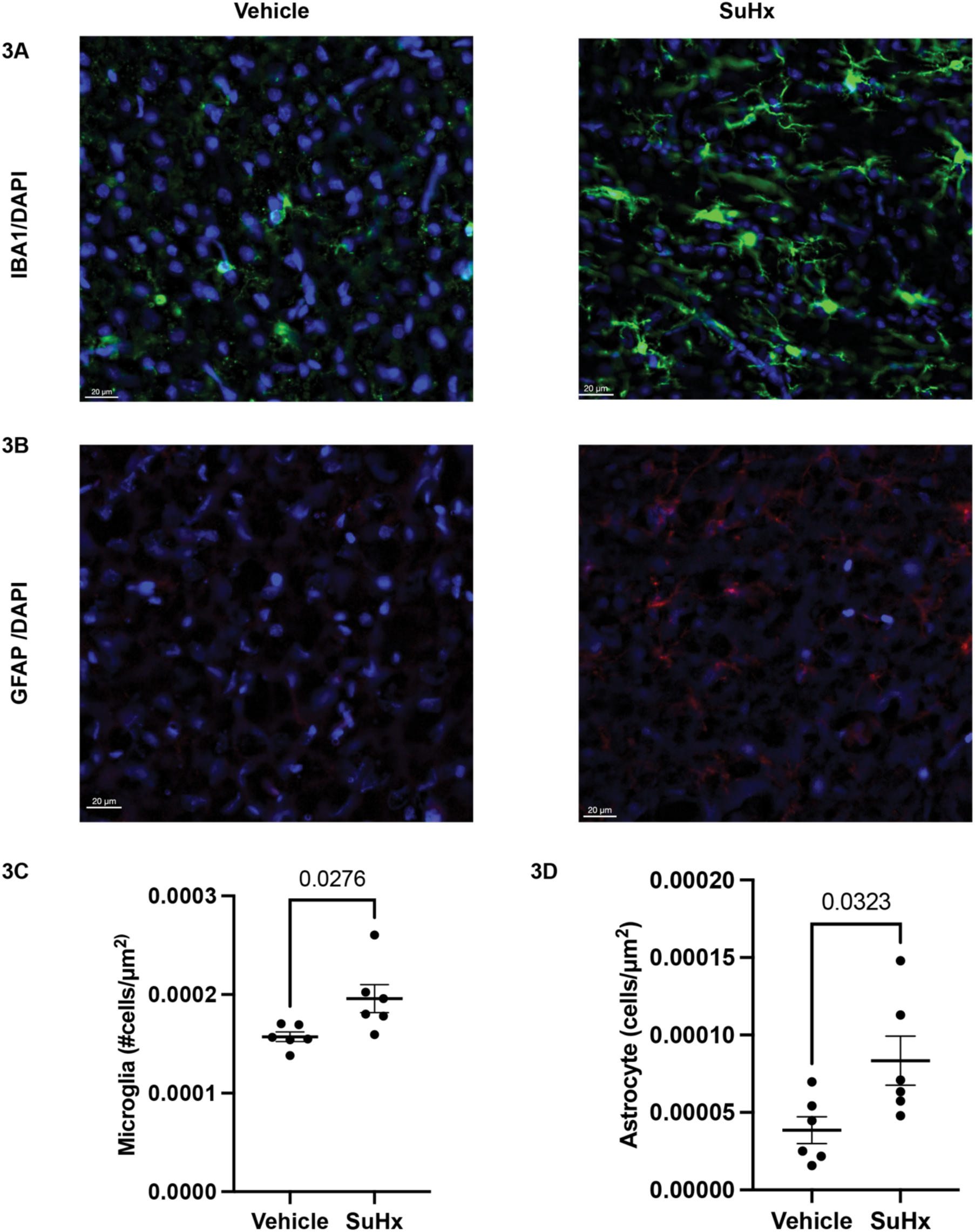
Sugen hypoxia increases cortical gliosis. Representative immunofluorescence images of fron­tal cortex microglia (IBA1/DAPI) **(3A)** and astrocytes (GFAP/DAPI) **(3B)** in rats exposed to sugen/hypoxia (SuHx) or vehicle. SuHx-treated rats (n = 6) exhibited a significantly increased microglial density (3.875 × 10^-5^ ± 1.504 × 10^-5^) compared to vehicle-treated controls (n = 6; p = 0.0276) (**3C**). Similarly, astrocyte densi­ty was elevated in SuHx-treated animals (4.487 × 10^-5^± 1.806 × 10^-5^) relative to controls (p = 0.0323) **(3D).** Data are presented as mean ± SEM and were analyzed using a two-tailed unpaired t-test.

**Figure 4.**
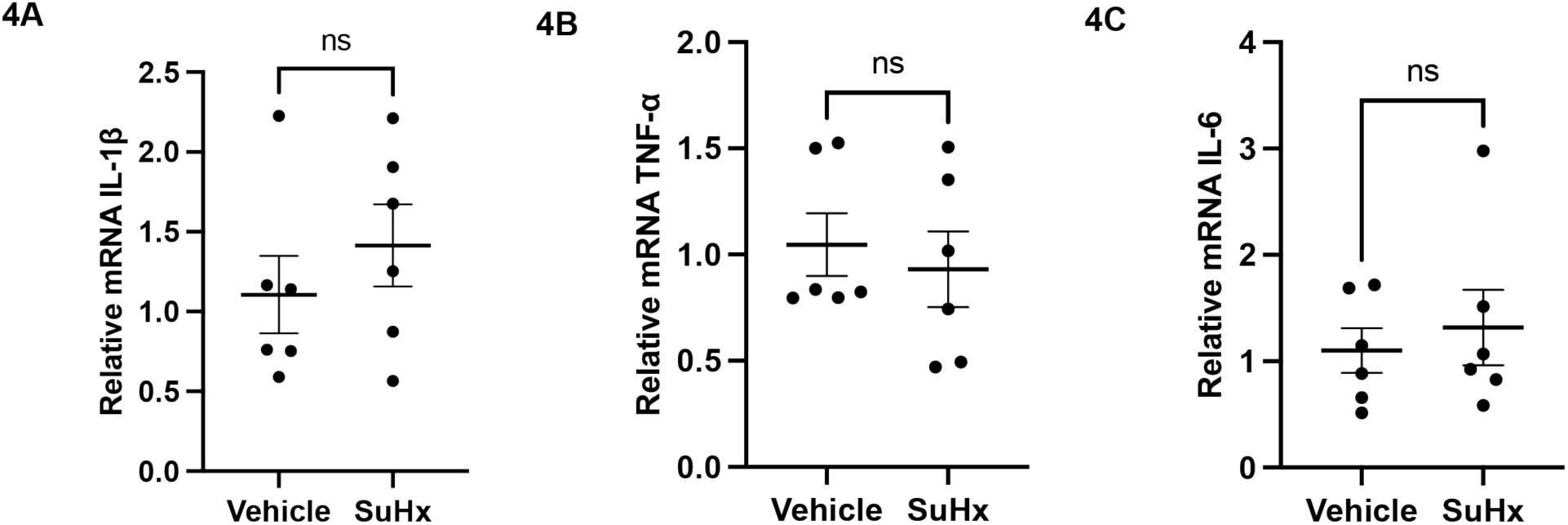
Sugen/hypoxia does not increase cortical cytokine expression. Quantitative PCR analysis of cortical cytokine mRNA expression in rats treated with sugen/hypoxia (SuHx; n = 6) or vehicle (n = 6) revealed no significant differences in IL-1 β **(4A),** TNF-α **(4B),** or IL-6 **(4C)** transcript levels. Data are presented as mean ± SEM and were analyzed using a two-tailed unpaired t-test.

### Experimental PH Induces Neuronal Loss and TDP-43 Proteinopathy

Neurodegeneration is characterized by the progressive loss of neurons and associated structural and functional decline^31^. A hallmark of several neurodegenerative diseases is TDP-43 proteinopathy, defined by the pathological mislocalization of TAR DNA-binding protein 43 (TDP-43) from the nucleus to the cytoplasm. While traditionally linked to conditions such as amyotrophic lateral sclerosis (ALS) and frontotemporal lobar degeneration (FTLD), TDP-43 pathology has also been identified in proinflammatory states, including traumatic brain injury and stroke^8,9,11,12^. In the SuHx model, we observed a significant reduction in cortical neuronal density compared to vehicle controls (n = 6), with a mean difference of –0.0001113 ± 2.937 × 10⁻⁵ cells/μm² (*p* = 0.0035). SuHx-treated animals also exhibited a significant increase in neuronal extranuclear TDP-43, with a mean difference of 0.2592 ± 0.0432 (*p* < 0.0001). Linear regression analysis revealed that right ventricular pressure (RVP) significantly predicted neuronal density (*R²* = 0.5825, *p* = 0.0039) and the proportion of neurons with extranuclear TDP-43 (*R²* = 0.4029, *p* = 0.0266) (**Figure 5**). Similarly, the Fulton index strongly predicted both neuronal density (*R²* = 0.6898, *p* = 0.0008) and extranuclear TDP-43 burden (*R²* = 0.6001, *p* = 0.0031) (data not shown).

**Figure 5.**
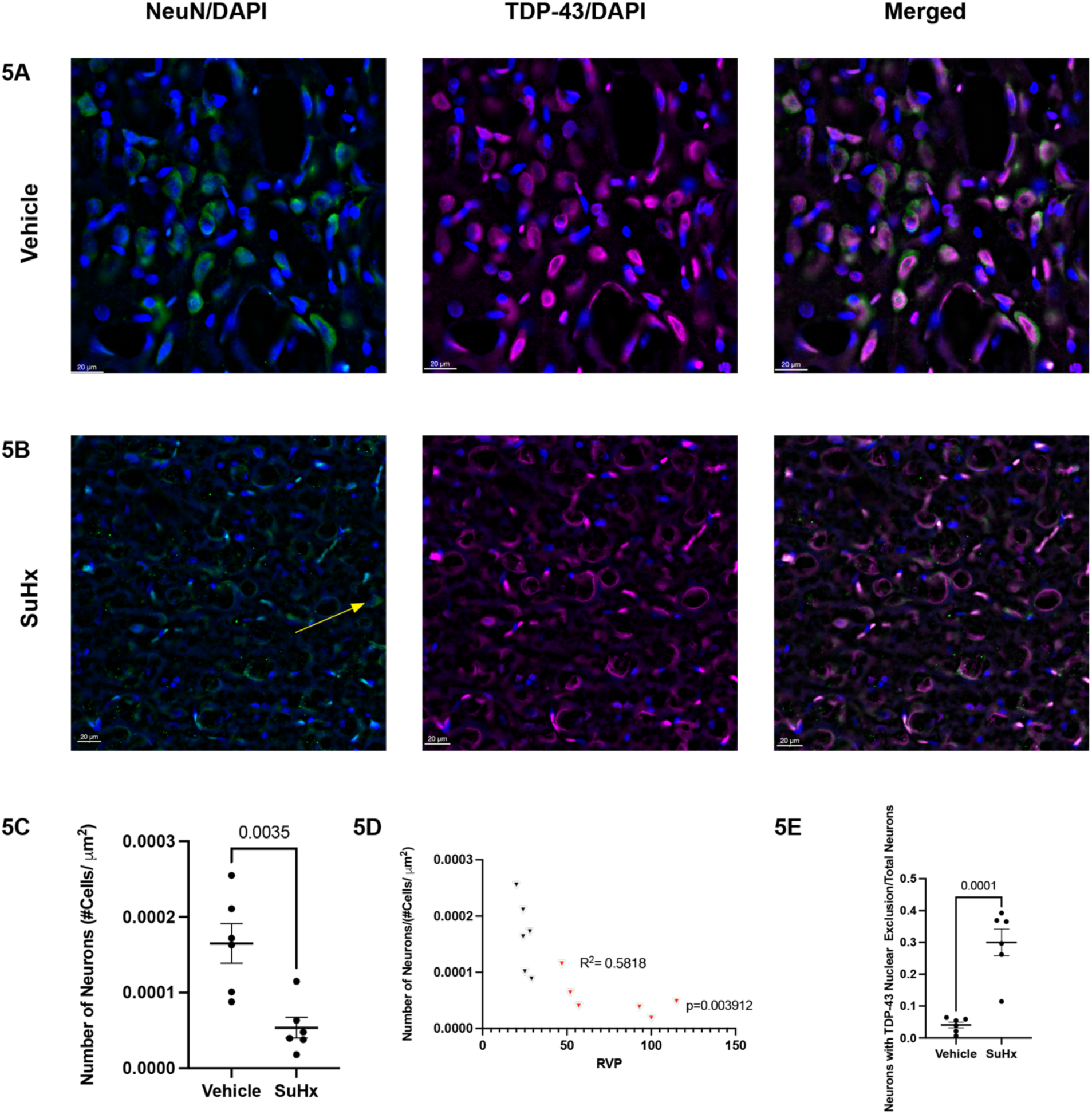
Sugen/hypoxia treatment reduces cortical neuronal density and induces neuronal TDP-43 mislocalization. Representative immunofluorescence images of cortical neurons stained for NeuN (neuronal marker) and DAPI (nuclear marker), TDP-43 and DAPI, and merged channels in rats exposed to vehicle at 4Ox magnification (**5A**) compared to sugen/hypoxia-treated rats (SuHx). The yellow arrow highlights dysmorphic neurons in SuHx-treated rats (**5B**). Quantification reveals a significant reduction in neuronal density in SuHx-treated rats (n=6) relative to vehicle controls (n=6), with a mean difference of-0.0001113 ± 2.937 x 10 ^5^ (p = 0.0035) (**5C**). Linear regression analysis indicates that right ventricu­lar pressure significantly predicts neuronal density, accounting for 58% of the variance (R^2^ = 0.5825, p = 0.0039). (**5D**). SuHx-treated animals also exhibit significantly increased neuronal TDP-43 nuclear exclusion compared to controls, with a mean difference of 0.2592 ± 0.04323 (p < 0.0001) (**5E**). Data are presented as mean ± SEM and analyzed using unpaired two-tailed t-tests unless otherwise specified.

In contrast, MCT-treated animals showed no significant change in neuronal density relative to controls (n = 6). However, MCT treatment was associated with a significant increase in neuronal extranuclear TDP-43 (mean difference: 0.2553 ± 0.0762, *p* = 0.0194) (**Supplemental Figure 5**). Notably, in the MCT group, neither RVP nor Fulton index correlated with neuronal density or extranuclear TDP-43.

### Clinical PH Induces Cortical TDP-43 Proteinopathy

To investigate the potential impact of pulmonary hypertension (PH) on neurodegeneration and TDP-43 proteinopathy in humans, we analyzed postmortem cortical brain tissue from individuals with PH (*n* = 4) and non-PH controls (*n* = 3). Quantitative analysis revealed no significant difference in cortical neuronal density between the two groups. However, neuronal extranuclear TDP-43 was significantly increased in PH subjects compared to controls, with a mean difference of 46.27 ± 12.96 (*p* = 0.0230) (**Figure 6**). Demographics available in supplemental table 2.

**Figure 6.**
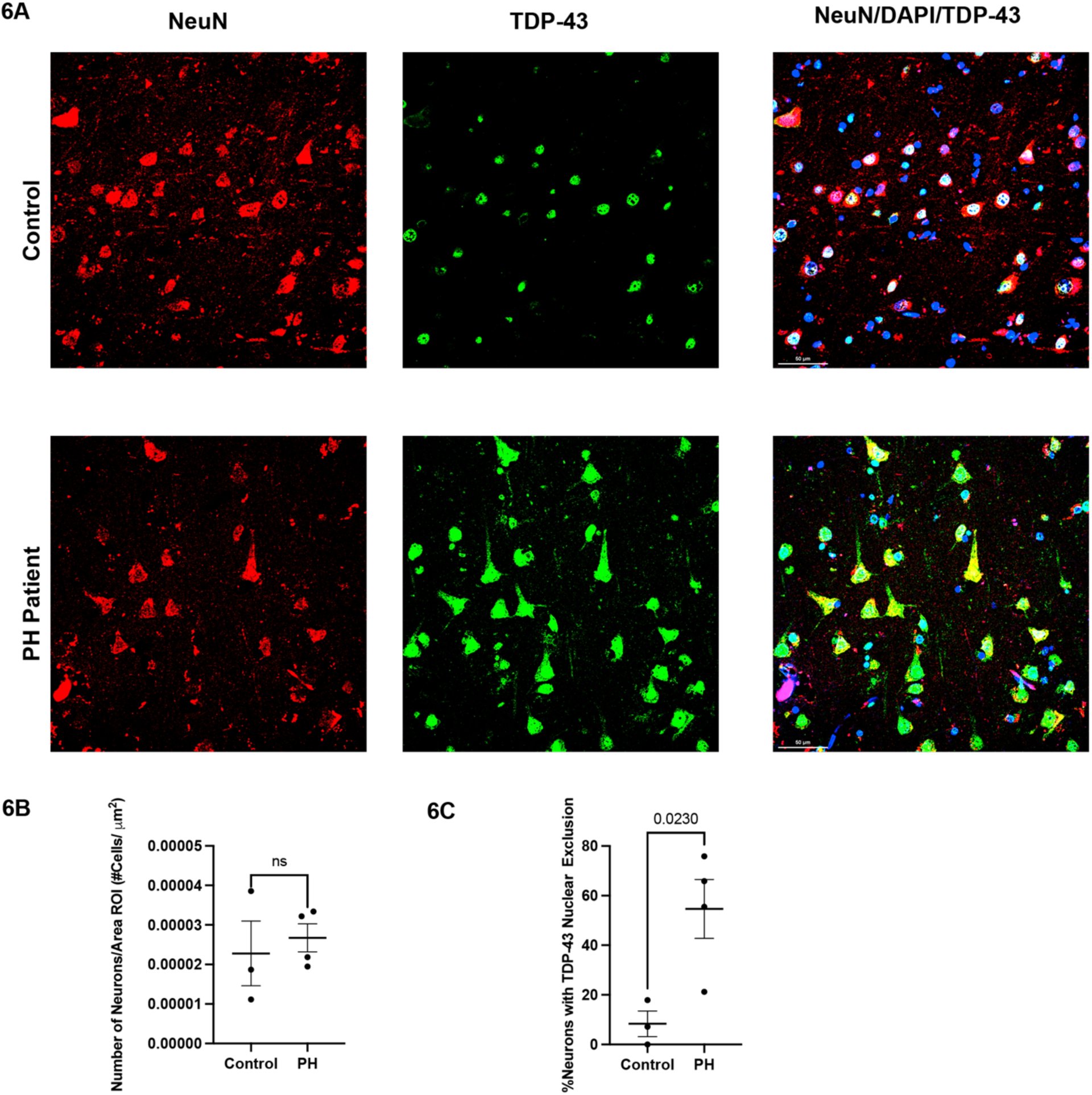
Clinical pulmonary hypertension is associated with increased neuronal TDP-43 mislocalization. Representative immunohistochemistry of frontal cortex tissue from patients with pulmonary hypertension (PH) and age-matched control subjects was used to assess neuronal density and TDP-43 localization **(6A).** Quantitative analysis revealed no significant difference in cortical neuronal density between PH and control groups (**6B**). However, PH subjects (n = 4) demonstrated significantly increased neurons with extranuclearTDP-43compared to controls (n = 3), with a mean difference of 46.27 ± 12.96 (p = 0.0230; **6C).** Scale bar = 50 µm. Data are presented as mean ± SEM and analyzed using a two-tailed unpaired t-test.

### PH Disproportionately Affects Expression of TDP-43 Regulated Genes in Pulmonary Artery Endothelial Cells (PAECs)

Emerging evidence suggests that TDP-43 is essential for endothelial cell function and RNA homeostasis^10,32^. This may represent a critical mechanistic link between TDP-43 proteinopathy and pulmonary hypertension (PH), implicating endothelial dysregulation as a contributor to the neurodegenerative processes observed in PH. To explore this hypothesis, we performed a secondary analysis of single-cell RNA sequencing data (GSE243193) from pulmonary artery endothelial cells^15^. We identified TDP-43 target transcripts using POSTAR3 CLIP-seq data and applied Fisher’s exact test to determine whether PH disproportionately dysregulates TDP-43–regulated genes^21^. Top downregulated genes were manually annotated using GeneCards^33^. This analysis revealed a significant overrepresentation of TDP-43–associated transcripts among the downregulated genes in PH (Fisher’s exact test: *p* < 2.2 × 10⁻¹⁶, OR: 0.193, 95% CI: 0.139-0.269), suggesting selective vulnerability of this RNA-regulatory pathway. Notably, many of the top downregulated TDP-43 targets are involved in RNA processing (EIF4A1, HNRNPH2), vesicular trafficking (SNX15, CHMP3, RAB4B), and inflammation (MIF) (**Figure 7**).

**Figure 7.**
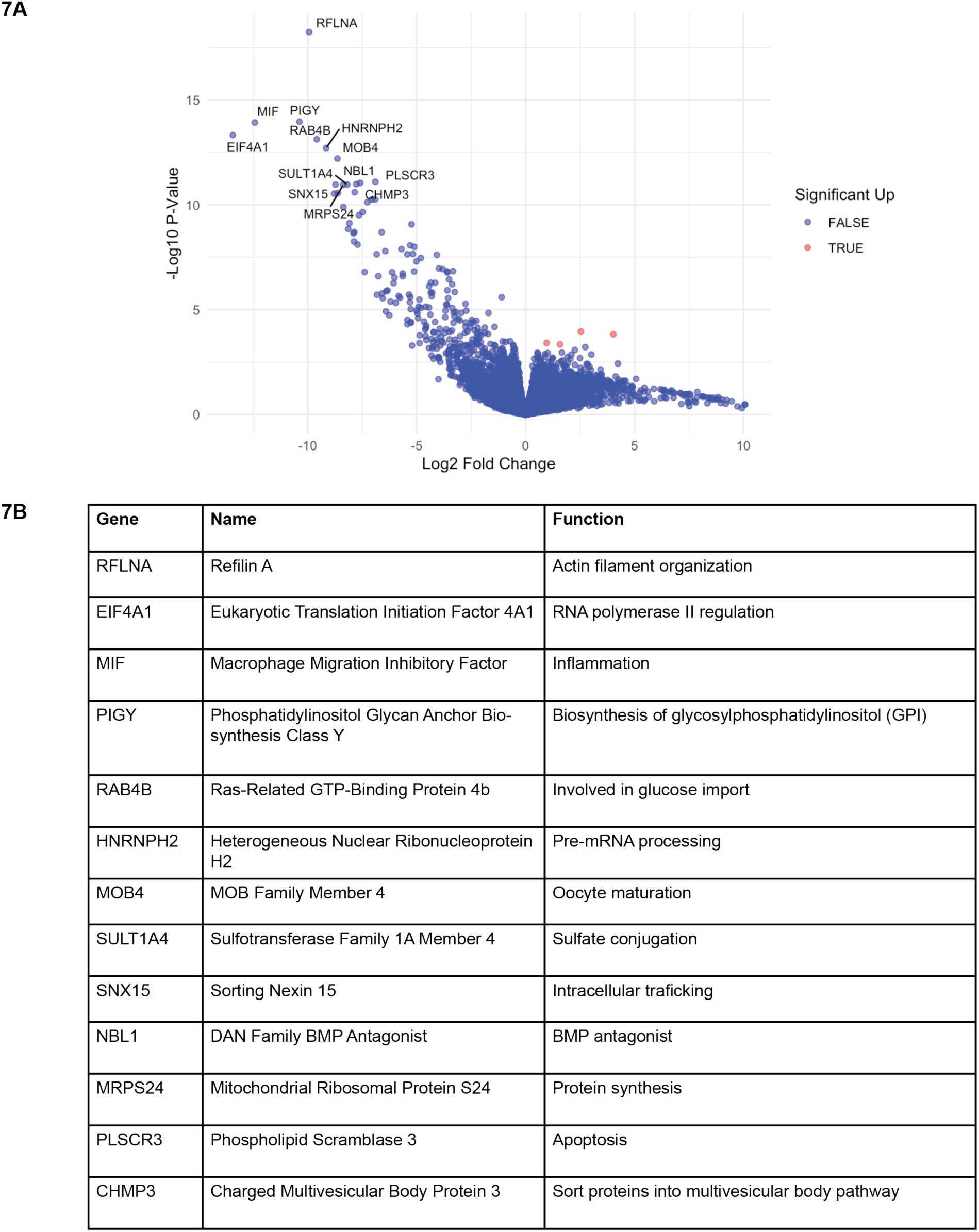
Differential expression of TDP-43 target genes in pulmonary artery endothelial cells from PH dataset. Volcano plot showing differential gene expression from single-cell RNA sequencing of pulmonary artery endothelial cells (PAECs) in PH (n=2O) vs control (n=3) (GSE243193). Fisher exact test confirmed that TDP-43-regulated genes were disproportionately downregulated in PH (p < 2.2 × 10^-16^, OR: 0.193, 95% Cl: 0.139-0.269) (**7A**). Manual annotation of top dyresgulated genes provided (**7B**).

## Discussion

Pulmonary hypertension (PH) is increasingly acknowledged as a systemic condition that affects not only the pulmonary vasculature and the right ventricle but also encompasses cognitive, affective, and neuroinflammatory consequences^2–4,7^. While prior clinical studies have documented neurocognitive symptoms in PH patients, the underlying pathophysiology linking pulmonary vascular disease to central nervous system (CNS) injury has remained poorly defined. Although previous clinical studies have documented neurocognitive symptoms in patients with PH, the pathophysiological mechanisms linking pulmonary vascular disease to central nervous system (CNS) injury remain inadequately defined. In this study, we present preclinical and translational evidence suggesting that gliosis and TDP-43 proteinopathy are contributing factors to the development of neurodegeneration in PH.

Our findings elucidate significant distinctions between the MCT and SuHx models of pulmonary hypertension (PH). Both models indicate a sex-related disparity in PH and vascular remodeling, aligning with existing literature^34^. As previously reported, the MCT induced a less severe PH phenotype in females, which resulted in right ventricular hypertrophy without elevated right ventricular pressure^34^. Notably, there are differences in central nervous system (CNS) findings between these models. Both models exhibit gliosis and TDP-43 proteinopathy; however, the more severe SuHx model also shows reduced neuronal density. The severity of PH correlates with TDP-43 proteinopathy and neuronal loss in the SuHx model, whereas in the MCT model, it predicts microglial activation. This suggests that CNS findings may occur on a spectrum, with milder forms of PH inducing neuroinflammation, while more severe PH leads to neuronal loss. The SuHx and MCT models of pulmonary hypertension exhibit distinct pathological mechanisms, which may have implications for extrapulmonary involvement, including the brain. The SuHx model combines VEGF receptor inhibition with chronic hypoxia to induce severe, angio-obliterative pulmonary vascular remodeling. This model simulates a progressive, endothelial-driven vasculopathy, which may potentially exacerbate blood-brain barrier (BBB) dysfunction and cerebral microvascular injury.^35^. Monocrotaline, in contrast, induces pulmonary hypertension through a singular, systemic endothelial toxin that preferentially damages the pulmonary vasculature via hepatic metabolism. Although effective in promoting vascular remodeling and right ventricular hypertrophy, monocrotaline is associated with systemic toxicity, including hepatic and renal injury, and may exert more variable effects on the cerebral vasculature. ^35^. Given the critical role of hypoxia in influencing neurovascular integrity and mitochondrial metabolism, the Sugen hypoxia model may more accurately represent brain-relevant systemic stressors, such as chronic cerebral hypoxemia. In contrast, the MCT model may complicate interpretation due to its extensive systemic toxicities. Our limited clinical data revealed an increase in TDP-43 proteinopathy among patients with PH compared to control subjects.

TDP-43, a nuclear RNA-binding protein with established roles in neurodegenerative diseases such as ALS and frontotemporal dementia, has recently been implicated in vascular and endothelial cell biology^10,32^ In these studies, the depletion of endothelial TDP-43 resulted in diminished expression of tight junction proteins, such as claudin-5, heightened permeability, and the activation of microglia and astrocytes.^36,37^. Although our studies on animal and human tissues concentrated on neuronal pathology, we hypothesized that endothelial dysfunction might contribute to PH-associated neuroinflammation through TDP-43–mediated post-transcriptional dysregulation. To investigate this hypothesis, we conducted a secondary analysis of single-cell RNA sequencing data from pulmonary artery endothelial cells (GSE243193), focusing on transcripts known to be regulated by TDP-43 using POSTAR3 CLIP-seq data^15,21^. This analysis demonstrated that PH disproportionately downregulates TDP-43 target genes in endothelial cells, with key affected transcripts involved in RNA metabolism (EIF4A1, HNRNPH2), vesicular trafficking (CHMP3, SNX15, RAB4B), mitochondrial homeostasis (MRPS24, PLSCR3), and inflammatory signaling (MIF)^33^. These findings support a model in which TDP-43–associated transcriptomic dysregulation may impair endothelial function in PH, potentially contributing to downstream neuroinflammation and blood-brain barrier vulnerability. While we did not directly assess blood-brain barrier (BBB) integrity in this study, the observed gene expression changes implicate pathways known to affect endothelial stability. Importantly, these data do not suggest that TDP-43 dysfunction is merely a consequence of chronic hypoxia or right heart failure. Rather, they propose that TDP-43 dysregulation is a shared molecular feature of both pulmonary vascular and neurologic injury in PH. This hypothesis aligns with recent literature demonstrating the role of TDP-43 in oxidative stress responses and RNA transport in both neurons and endothelial cells^9,10,32,38^.

Our study has several limitations. Firstly, although the neuroinflammatory and degenerative alterations were pronounced in both animal and human PH brain tissue, we did not directly demonstrate functional impairments, such as behavioral or cognitive deficits. Further work is needed to phenotype cognitive impairment in PH animal models. While an open field test is beneficial for identifying locomotion, anxiety, and sickness behavior in animal models, it may not effectively simulate the loss of executive function observed in frontotemporal lobar degeneration (FTLD), which is also challenging to differentiate from motor dysfunction in a clinical context ^39^. Next, quantitative PCR analysis did not reveal a significant difference in proinflammatory cytokine mRNA levels, indicating that further investigation is required to determine the timing of neuroinflammation in relation to the onset of pulmonary hypertension. An additional limitation is the use of a limited data tissue biobank utilized to investigate TDP-43 in PH patients. This dataset does not include information regarding PH duration or severity which could influence the results. In addition, while our transcriptomic analysis indicates endothelial contributions to the TDP-43 signature, further research is necessary to validate these targets at the protein level and to assess causality using endothelial-specific TDP-43 models. Lastly, our study was not powered to assess sex as a biological variable which may be an important direction for future studies. In sum, the temporal sequence of neuroinflammation, TDP-43 mislocalization, endothelial dysfunction, and CNS injury remains to be elucidated.

Despite these limitations, our findings establish a foundational basis for a novel conceptual model of PH as a multisystem disorder characterized by mechanistic links to RNA dysregulation and neurodegeneration. Targeting TDP-43 or its downstream pathways may present therapeutic opportunities to mitigate both vascular and neurological complications in this population. Future research focused on elucidating the cell-specific roles of TDP-43 within the lung-brain axis may yield further insights into the systemic nature of pulmonary vascular disease.

### Conclusion

This study identifies TDP-43 dysregulation as a potential mechanistic link between pulmonary hypertension and central nervous system injury. Through the integration of animal models, human brain tissue, and endothelial transcriptomic analysis, we demonstrate that PH is associated with neuroinflammatory and neurodegenerative changes, alongside selective downregulation of TDP-43–regulated genes in pulmonary artery endothelial cells. These findings support a novel model in which TDP-43 contributes to multisystem dysfunction in PH, bridging vascular and neurologic pathology. Elucidating the role of TDP-43 across organ systems may open new avenues for targeted therapies aimed at mitigating both cardiopulmonary and cognitive complications of PH.

## Author Contributions

PD-Designed and performed experiments, data analysis, manuscript preparation, AL-Performed and interpreted cell culture experiments, MG-Performed and interpreted cell culture experiments, SB-Performed antibody validations, immunostaining and microscopy, AR-Performed antibody validations, immunostaining and microscopy, AF-Managed animal colonies, TC-Performed and interpreted right heart catheterization, CL-Assisted with design of immunohistochemistry experiments, SM-Experimental design, technical expertise, SU-

## Abbreviations

(PH): Pulmonary Hypertension
(TDP-43): TAR DNA-binding protein 43
(ALS): Amyotrophic Lateral Sclerosis
(FTLD): Frontotemporal Lobar Degeneration
(PAEC): Pulmonary artery endothelial cell
(CNS): Central nervous system
(DMSO): Dimethyl sulfoxide
(MCT): Monocrotaline
(RVP): Right Ventricular Systolic Pressure
(PBST): Phosphate Buffered Saline-Triton X
(Iba1): Ionized calcium-binding adaptor molecule 1
(GFAP): Glial Fibrillary Acidic Protein
(NeuN): Neuron-Specific Nuclear Protein
(PCR): Polymerase Chain Reaction
(SuHx): Sugen/Hypoxia
(DEGs): Differentially expressed genes
(FDR): False discovery rate

## Acknowledgements

This work was supported by the Behavioral Phenotyping Core (BPC) at the Indiana University School of Medicine.

## Figures

**Supplemental Table 1.** Antibodies, dilution, manufacturer, and product ID utilized in immunohistochemistry experiments.

| Antibody | Dilution | Manufacturer | Product ID |
| --- | --- | --- | --- |
| Ionized calcium-binding adaptor molecule 1 (Iba1) | 1:500 | Synaptic Systems | #234009 |
| Glial Fibrillary Acidic Protein (GFAP) | 1:1000 | Invitrogen | #13-0300 |
| TDP-43 | 1:500 | Proteintech | #10782-2 |
| Neuron-Specific Nuclear Protein (NeuN) | 1:1000 | Antibodies.com | #A270544 |

**Supplemental Table 2.** Pulmonary hypertension subject demographics. Abbreviations include; unavailable (UA), Chronic Thromboembolic Pulmonary Hypertension (CTEPH), Portal pulmonary hypertension (PoPH), end stage renal disease (ESRD), non-alcoholic steatohepatitis (NASH), alcohol use disorder (AUD), diabetes mellitis 2 (DM2), autoimmune thrombocytopenic purpura (ITP), deep venous thrombosis (DVT).

| Age | Sex | PH Etiology | PH Severity (RVP) | Comorbidities |
| --- | --- | --- | --- | --- |
| 69 | F | Control | UA | UA |
| 27 | F | Control | UA | UA |
| 64 | F | Control | UA | UA |
| 83 | M | CTEPH | UA | UA |
| 69 | M | PoPH | 43 mmHg | ESRD, NASH cirrhosis |
| 47 | M | PoPH | 52 mmHg | AUD, cirrhosis, DM2, hepatic encephalopathy |
| 43 | F | SLE-PH | 93 mmHg | stroke, ITP, DVT |

**Supplemental Figure 1.**
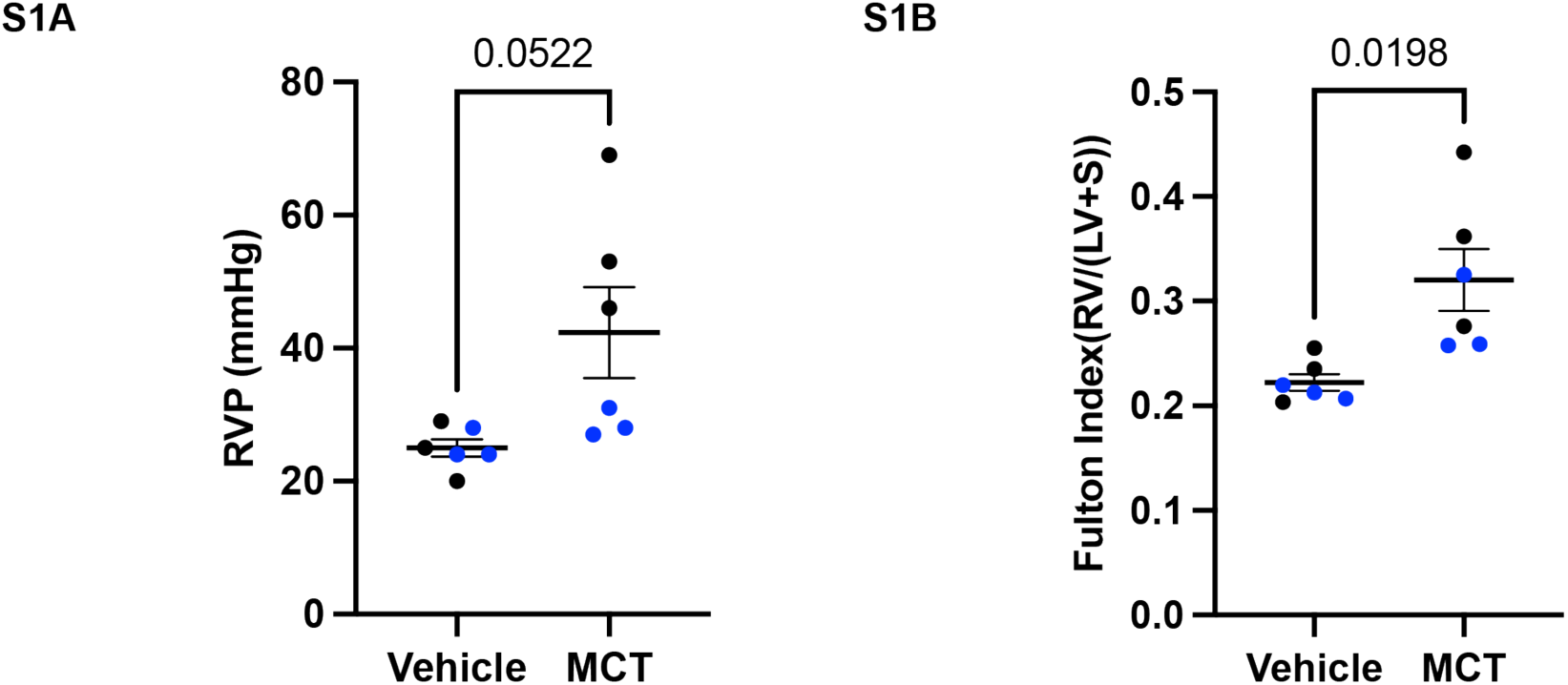
Monocrotaline induces vascular remodeling. Rats treated with monocrotaline (MCT; n = 6) trend to an increased right ventricular pressure (RVP), with a mean difference of 17.33 ± 6.98 mmHg compared to vehicle controls (n = 6; p = 0.0522) **(S1A).** MCT-treated rats demonstrate right ventricular hypertrophy, with a mean Fulton Index difference of 0.09815 ± 0.03066 compared to controls (p = 0.0198) **(S1B).** Data analyzed using a two-tailed unpaired t-test and are presented as mean± SEM. Female animals depicted in blue.

**Supplemental Figure 2.**
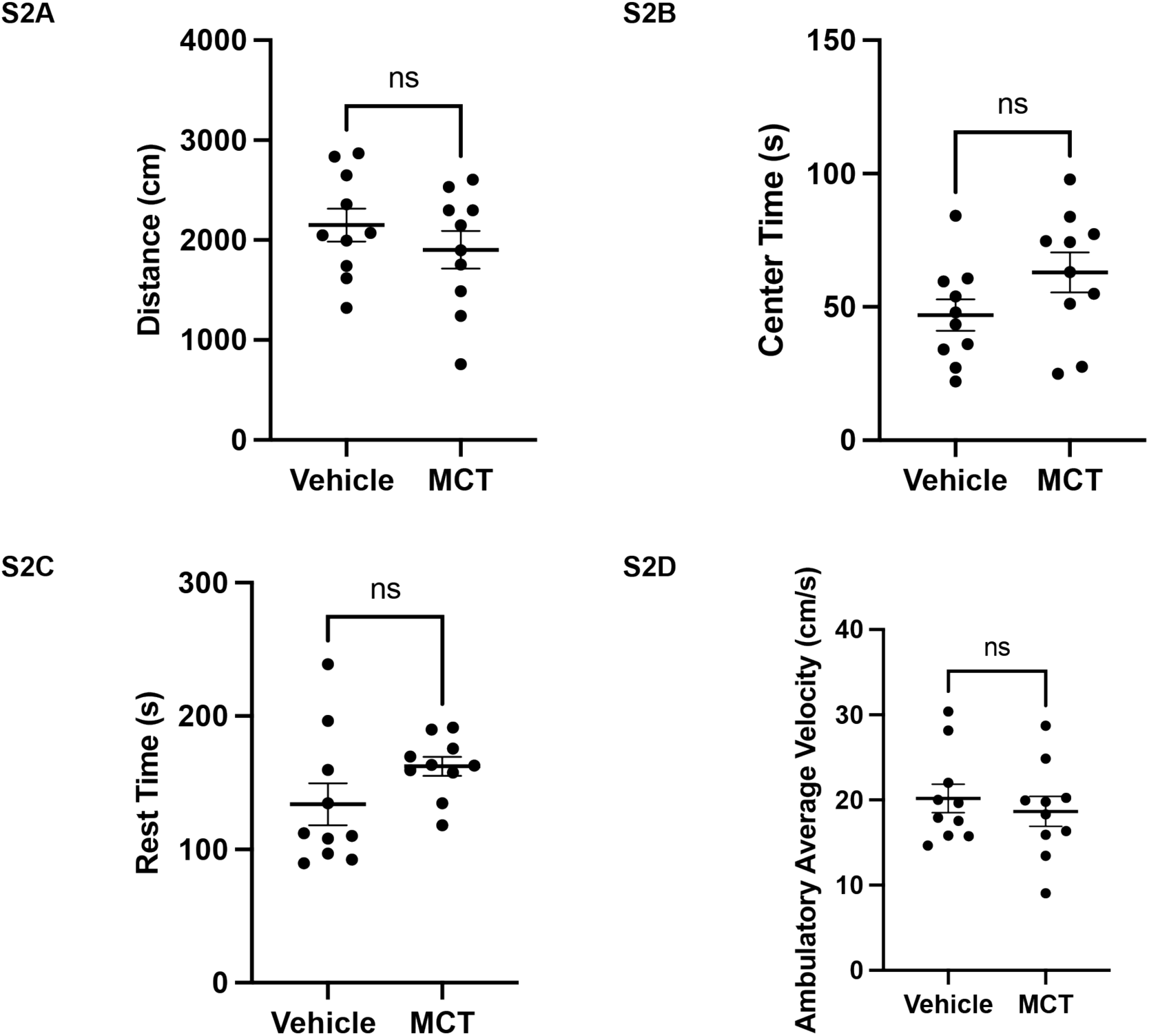
Monocrotaline does not induce sickness behavior in rats undergoing open field test. Rats exposed to Monocrotaline (MCT, n = 10) did not exhibit significant differences in behavioral parameters compared to vehicle-treated controls (n = 10). Specifically, there were no significant changes in total distance traveled **(S2A),** time spent in the center of the open field test **(S2B),** time spent at rest **(S2C),** or average ambulatory velocity **(S2D).** Data were analyzed using an unpaired two-tailed t-test and are presented as mean ± SEM.

**Supplemental Figure 3.**
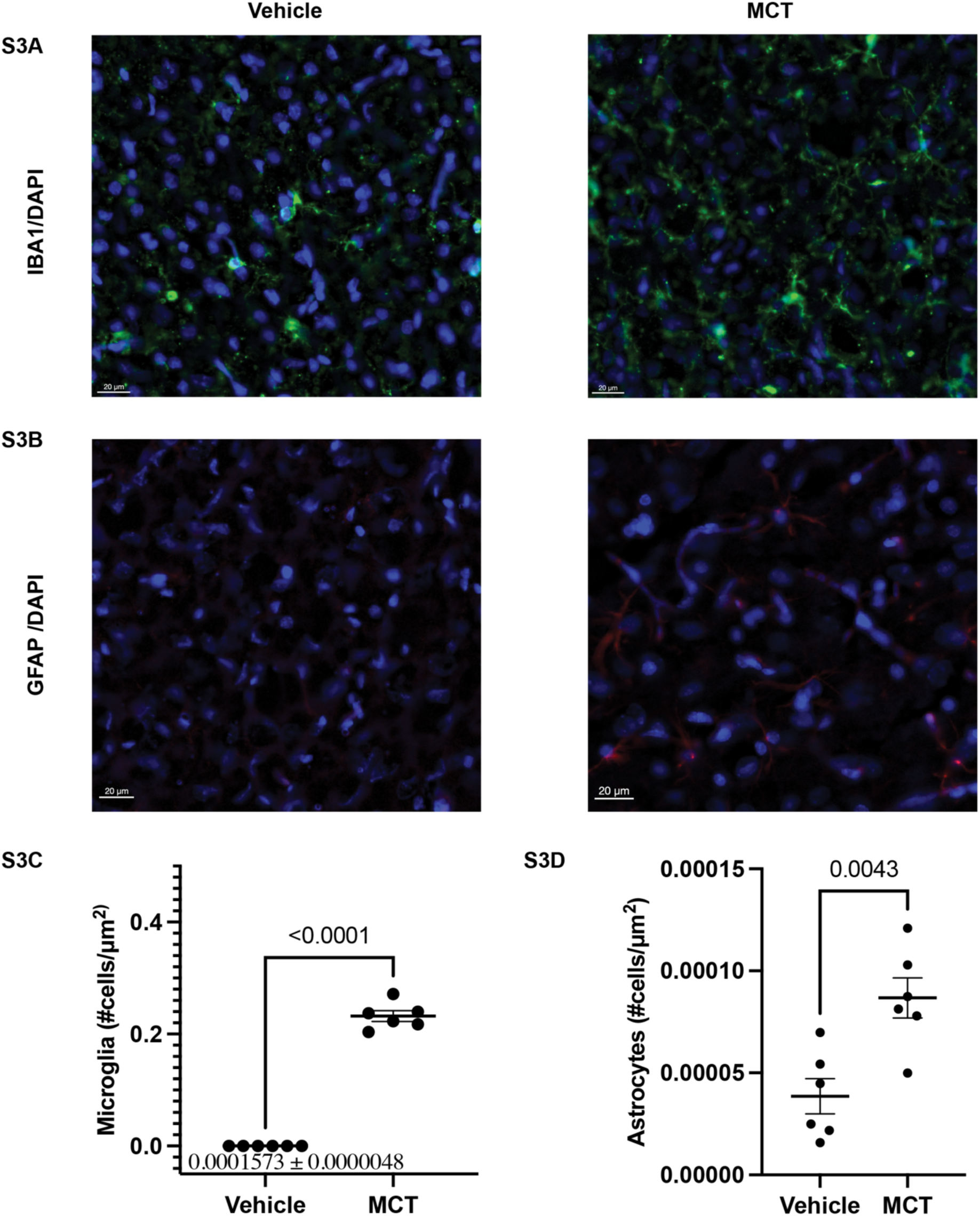
Monocrotaline increases cortical gliosis. Representative immunofluorescence images of frontal cortex microglia (IBA1/DAPI) (**S3A**) and astrocytes (GFAD/DAPI) (**S3B**) in rats exposed to monocrotaline (MCT) or vehicle. MCT-treated rats (n = 6) exhibited a significantly increased microglial density (0.2320 ± 0.009541) compared to vehicle-treated controls (n = 6; p < 0.0001). (**S3C**). Similarly, astrocyte density was elevated in MCT-treated animals (4.822 X 10^-5^) realtice to controls (p = 0.0043) (**S3D**). Data are presented as mean ± SEM and were analysed using a two-tailed unpaired t-test. Error bars are shown for all groups; however, the control group's errror bars are not visible at this scale due to extremely low variance (mean = 0.0001573, DEM = 0.00000048) and large difference from the MCT group. A text label is included to indicated the control mean ± SEM.

**Supplemental Figure 4.**
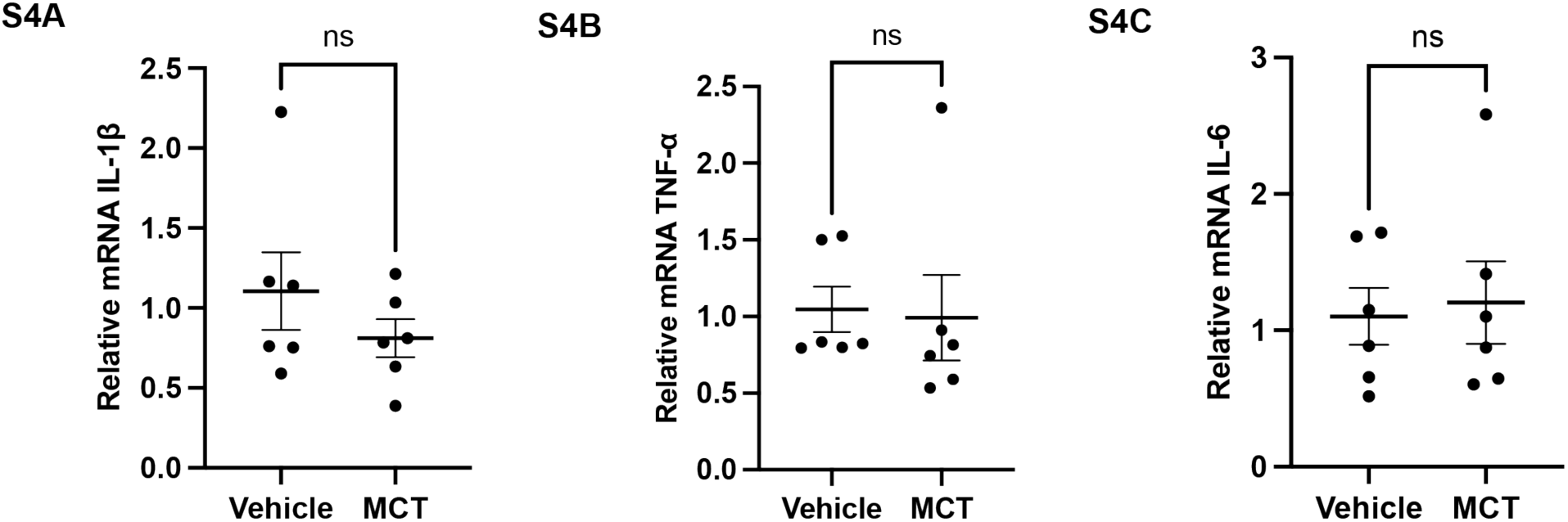
Monocrotaline does not increase cortical cytokine expression. Quantitative PCR analysis of cortical cytokine mRNA expression in rats treated with monocrotaline (MCT; n = 6) or vehicle (n = 6) revealed no significant differences in IL-1 β **(S4A),** TNF-α **(S4B),** or IL-6 **(S4C)** transcript levels. Data are presented as mean ± SEM and were analyzed using a two-tailed unpaired t-test.

**Supplemental Figure 5.**
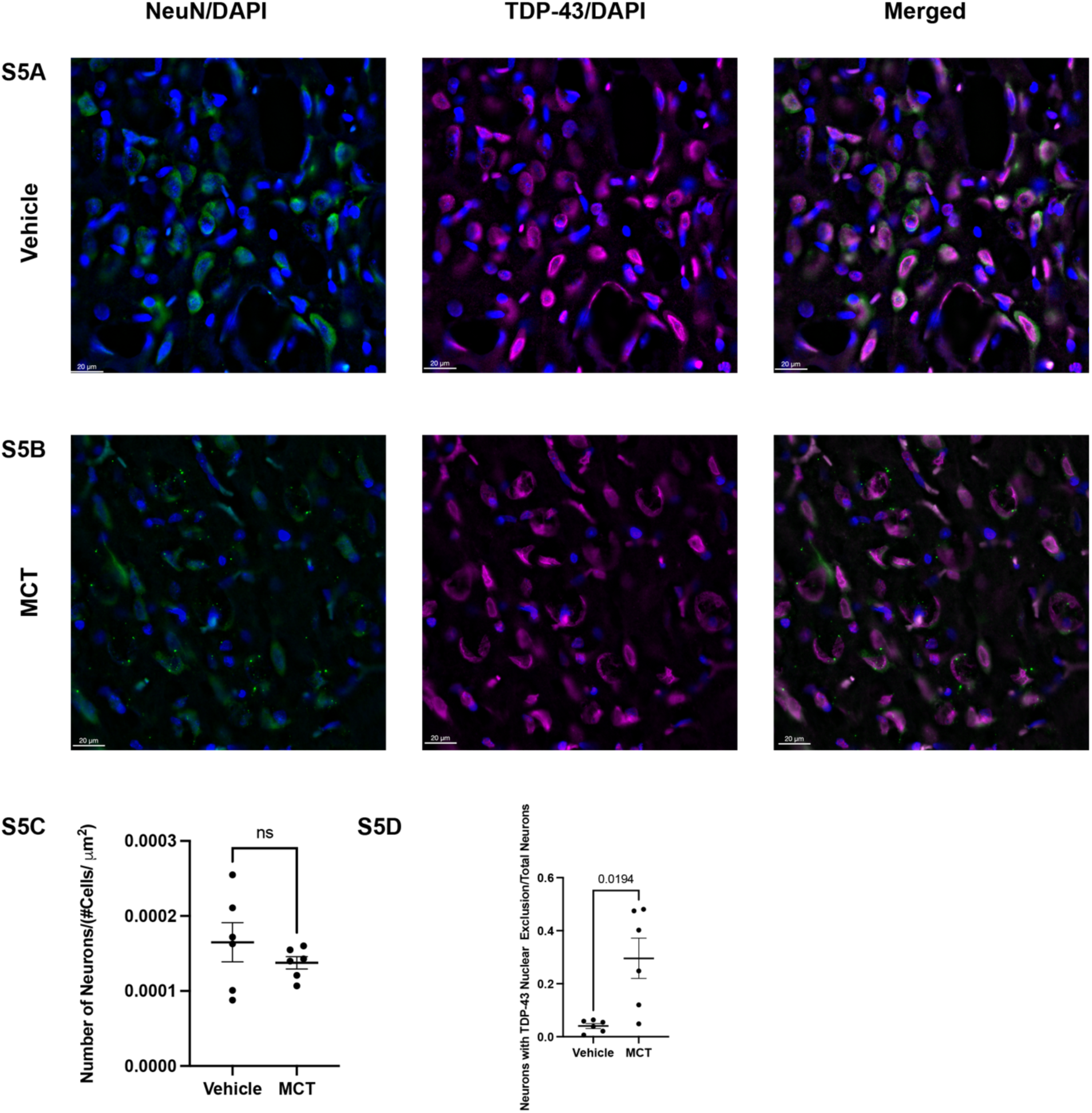
Monocrotaline treatment reduces cortical neuronal density and induces neuronal TDP-43 mislocalization. Representative immunofluorescence images of cortical neurons stained for NeuN (neuronal marker) and DAPI (nuclear marker), TDP-43 and DAPI, and merged channels in rats exposed to vehicle at 4Ox magnification **(S5A)** compared to monocrotaline-treated rats (MCT) **(S5B).** Quantification reveals no change in cortical neuronal density in MCT-treated rats (n=6) compared to vehicle controls (n=6) **(S5C).** MCT-treated animals exhibit significantly increased neuronal nuclear TDP-43 exclusion compared to controls, with a mean difference of 0.2553 ± 0.07623 (p = 0.0194) **(S5D).** Data are presented as mean ± SEM and analyzed using unpaired two-tailed t-test.

